# GRNPred: A Multimodal Graph Transformer with Masked Gene Expression Pretraining for Gene Regulatory Network Inference

**DOI:** 10.64898/2026.04.26.720917

**Authors:** Thong Nguyen, Akshata Hegde, Jialin Cheng

## Abstract

Gene regulatory network (GRN) inference is a fundamental problem in systems biology, aiming to identify transcription factor (TF)–target gene interactions from high-dimensional gene expression data. Accurate GRN reconstruction remains challenging due to limited labeled regulatory data, severe class imbalance, and the complex, nonlinear nature of transcriptional regulation. Here, we introduce GRNPred, a multimodal graph transformer framework for robust GRN inference that integrates gene expression, functional annotations, semantic gene descriptions, regulatory binding motif priors, and gene co-expression network topology. GRNPred follows a two-stage training strategy. In the first self-supervised pretraining phase, a graph transformer encoder is trained on TF-centered gene co-expression subgraphs using masked gene-expression reconstruction, enabling the model to learn transcriptional context from unlabeled data. In the second supervised fine-tuning stage, the pretrained encoder is finetuned for supervised TF–target edge prediction using available regulatory annotations. Transformer-based attention allows GRNPred to capture long-range and context-dependent regulatory interactions that are difficult to model with conventional graph neural networks. Extensive evaluation across 7 benchmark datasets and 3 regulatory network constructions demonstrates that GRNPred consistently outperforms state-of-the-art GRN inference methods, achieving AUROC scores of up to 0.94 and AUPRC scores of up to 0.93, while maintaining strong robustness across diverse biological contexts.

## 1 Introduction

Gene regulatory networks (GRNs) provide a compact representation of the gene regulatory logic governing cellular behavior by encoding interactions between transcription factors (TFs) and their target genes, which plays a central role in controlling cell identity, function and development. Consequently, accurate GRN inference from high-throughput data has become a core objective in computational systems biology and regulatory genomics.

Despite rapid advances in RNA sequencing (RNA-seq) technologies, inferring GRNs from gene-expression profiles remains challenging due to the high dimensionality of RNA-seq data, limited sample sizes, pervasive noise, and the complex, nonlinear nature of transcriptional regulation. Traditional GRN inference methods—including information-theoretic approaches such as ARACNe [1], tree-based ensemble models such as GENIE3 [2], and boosting-based variants like GRNBoost [3]—have achieved notable success but rely heavily on assumptions of pairwise dependence or static regulatory relationships. As a result, these methods often struggle to model higher-order interactions, cell-type-specific regulation, and long-range dependencies in large-scale gene networks.

More recently, machine-learning and deep-learning approaches have shown promise in capturing complex gene-regulatory patterns from expression data. Neural-network-based models can flexibly approximate nonlinear relationships and integrate heterogeneous data sources. Among these, graph neural networks (GNNs) have gained particular attention because they naturally model biological systems as graphs, enabling information propagation between genes via known or inferred interactions. GNNs have demonstrated strong capability to learn structured dependencies between nodes in graphs [4, 5]. In the context of GRNs, they allow each gene to aggregate information from its neighbors, thereby capturing regulatory cascades and modular organization [6].

However, conventional message-passing GNNs are limited by their local receptive fields and tend to oversmooth node representations as network depth increases [7]. Graph transformer architectures address these limitations by leveraging attention mechanisms to model global context and adaptive neighborhood interactions [8, 9]. By allowing each node to attend selectively to other nodes, graph transformers can capture long-range regulatory dependencies and context-specific interactions that are difficult to express with traditional GNNs. Recent studies have demonstrated the effectiveness of transformer-based graph models in modeling biological interaction networks, and systems-level prediction tasks have been significantly advanced by recent developments in graph neural networks and transformer-based architectures [10, 9, 11].

A further challenge in GRN inference is the scarcity and incompleteness of experimentally validated regulatory edges (ground truth labels). Gold-standard GRNs are often derived from ChIP-seq or perturbation experiments, which are expensive, context-specific, and unavailable for many cell types. As a result, supervised learning approaches risk overfitting to limited annotations and may not be generalized across datasets. Self-supervised learning provides a principled solution to this challenge by enabling models to learn informative representations directly from unlabeled data. Masked-reconstruction objectives, initially popularized in natural language processing via BERT [12], have been successfully adapted to graphs, as exemplified by GraphMAE and related frameworks [13]. These methods encourage models to capture intrinsic structure by predicting missing node features or masked subgraphs without relying on task-specific labels.

In this work, we introduce **GRNPred**, a graph transformer-based framework for GRN inference that unifies self-supervised representation learning with supervised regulatory edge prediction. GRN-Pred employs a two-stage learning strategy (Figure 1). In the pretraining stage (Figure 1a), a graph transformer encoder learns gene representations by reconstructing masked gene expression signals, enabling it to capture global expression landscapes and co-expression structure. In the fine-tuning stage (Figure 1b), the pretrained encoder is adapted for TF–target regulatory edges using available regulatory annotations (labels). This design allows GRNPred to leverage large-scale unlabeled gene expression data while remaining sensitive to biologically grounded regulatory constraints.

**Figure 1:**
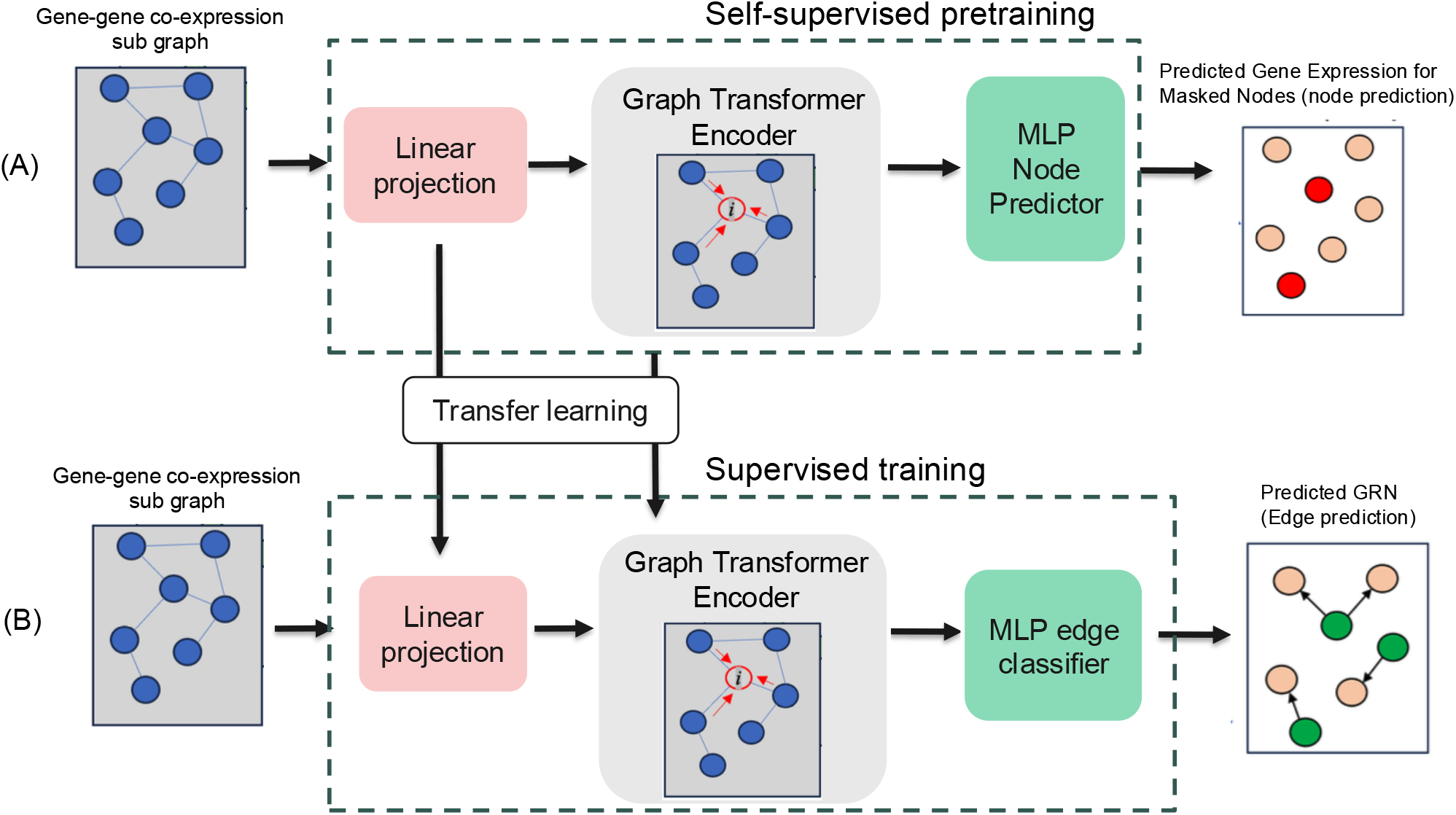
Overview of GRNPred. (A) In the self-supervised pretraining phase, GRNPred is trained to predict masked gene expression values (node prediction); (B) In the supervised fine-tuning phase, GRNPred is trained to predict TF-gene regulatory relationships (edge prediction). The initial weights for the fine-tuning phase are transferred from the weights learned in the pretraining phase.

By combining transformer-based global attention with self-supervised pretraining, GRNPred addresses several limitations of prior GRN inference methods. The model captures nonlinear, context-dependent regulation, learns robust representations under label scarcity. These properties make GRNPred a scalable and flexible framework for reconstructing transcriptional regulatory networks from high-dimensional gene-expression data.

## 2 Results

### 2.1 GRNPred overview

GRNPred is a graph transformer architecture designed to integrate heterogeneous biological signals into a unified representation of regulatory structure (Figure 1). The model operates on transcription factor (TF)–centered gene co-expression subgraphs, each capturing the local regulatory neighborhood surrounding a TF and its candidate target genes. Transformer-based graph architectures have demonstrated strong capacity for modeling long-range dependencies and complex relational patterns in biological networks [14]. Each input subgraph for GRNPred is constructed as a 2-hop TF-centered neighborhood [**?**] in a full gene co-expression network, ensuring coverage of both direct regulatory interactions and second-order contextual genes. This subgraph-centric formulation is closely aligned with recent transformer-based GRN inference frameworks, such as GRNFormer, which similarly emphasize TF-centered graph representations to enhance regulatory signal localization and predictive accuracy. By focusing computation on biologically meaningful regulatory units, this design balances expressiveness with scalability across large gene sets.

Each node in a GRNPred subgraph is represented using a multimodal feature vector composed of three components: (i) gene-expression values that capture transcriptional activity, (ii) Gene Ontology (GO) term embeddings summarizing functional annotations [15], and (iii) semantic embeddings derived from descriptive gene text using a LLaMA-based large language model [16], providing complementary contextual information. These heterogeneous components are concatenated to form the raw node representation and then passed through a multilayer perceptron (MLP) that maps them into a unified feature space. This projection mechanism follows the same design principles used in transformer-based architectures such as BERT [12] and GraphMAE [13], where input modalities are standardized into a common representation before attention-based processing.

The edges of each subgraph are enhanced with regulatory features, including regulatory motif-based binding scores and TF binding probabilities[13]. These edge features are processed through a dedicated MLP to generate dense edge embeddings compatible with the graph transformer layers.

At the core of GRNPred is a multi-layer graph transformer encoder. Each layer performs attention-based message passing across the TF subgraph, enabling the model to integrate long-range regulatory dependencies that extend beyond local neighborhoods, a capability characteristic of graph transformer models [9, 10]. A learned positional embedding is added to each node, allowing the network to incorporate structural context analogous to positional encoding in natural language transformers [12]. Across successive layers, the encoder synthesizes expression signals, functional annotations, semantic embeddings, and sequence-derived regulatory evidence into a unified latent representation.

The final output of the encoder is a set of high-dimensional contextual embeddings that summarize the regulatory potential of each TF–gene pair within the subgraph. These embeddings provide the foundation for downstream tasks such as edge prediction and expression-driven target characterization. By tightly integrating multimodal biological information with transformer-based graph modeling, GRNPred offers a flexible and expressive framework for uncovering gene regulatory structure.

Building on these embeddings, GRNPred employs a multilayer perceptron (MLP) edge classifier to compute final edge-level regulatory scores. For each TF–gene pair, the MLP classifier ingests a composite feature vector consisting of both node embeddings and interaction features such as element-wise products and absolute differences, consistent with established neural link-prediction methods [17].

This vector is passed through a series of linear transformations, normalization layers, and nonlinear activations to produce a scalar prediction reflecting the likelihood of a true regulatory interaction. This design allows GRNPred to effectively integrate node context, pairwise relationships, and learned nonlinear transformations into a discriminative edge-scoring function

### 2.2 Accurate Inference of GRNs across regulatory contexts

To rigorously evaluate the accuracy, stability, and biological relevance of GRNPred, we conducted a comprehensive benchmarking study across seven human cell types, each paired with three ground-truth regulatory networks. The datasets encompass human embryonic stem cells (hESC), human hepatocytes (hHep), dendritic cells (mDC), and four hematopoietic stem and progenitor cell subtypes (mHSC-L, mHSC-GM, mHSC-E, and mHSC). For each cell type, regulatory edges were evaluated against three distinct ground-truth networks commonly used in GRN benchmarking: (1) non–cell-type-specific ChIP-seq networks, (2) STRING-derived interaction networks[18], and (3) cell-type-specific ChIP-seq networks [19]. This arrangement provides a diverse spectrum of regulatory evidence ranging from direct TF-binding events to regulatory protein-protein interactions in STRING.

GRNPred is trained and evaluated independently on each dataset for gene regulatory network (GRN) inference. For every cell type and each TF–gene graph variant (tf500 and tf1000), the positive regulatory edges are split into training, validation, and test sets using a strict 70/15/15 ratio. Here, tf500 and tf1000 denote graphs built from the union of the curated TF list and the top 500 or top 1000 most variable genes (by expression variability) in the dataset, respectively, which controls the size and complexity of the candidate regulatory network. The subsets were prepared according to BEELINE’s standard data preprocessing and evaluation protocols.

Importantly, TFs are partitioned such that each TF appears in exactly one of the training, validation, or test sets, ensuring that TFs are disjoint and unique across splits and preventing information leakage between evaluation stages. The model is trained exclusively on the training split and tuned using the validation split, while the held-out test edges are used solely for final evaluation. This design ensures that GRNPred learns high-fidelity regulatory signals within each dataset while maintaining consistent and unbiased class distributions across all evaluation stages.

To obtain statistically robust and variance-stable performance estimates, we performed 100 rounds of bootstrap evaluation for every model– dataset–network combination [20, **?**] (Figure 2). Bootstrap-based testing is particularly important in GRN inference settings, where the number of negative TF–target pairs vastly exceeds the number of experimentally validated positive regulatory edges, leading to severe class imbalance. Under such conditions, evaluation on the full test set can yield misleading or unstable metrics dominated by negative examples.

**Figure 2:**
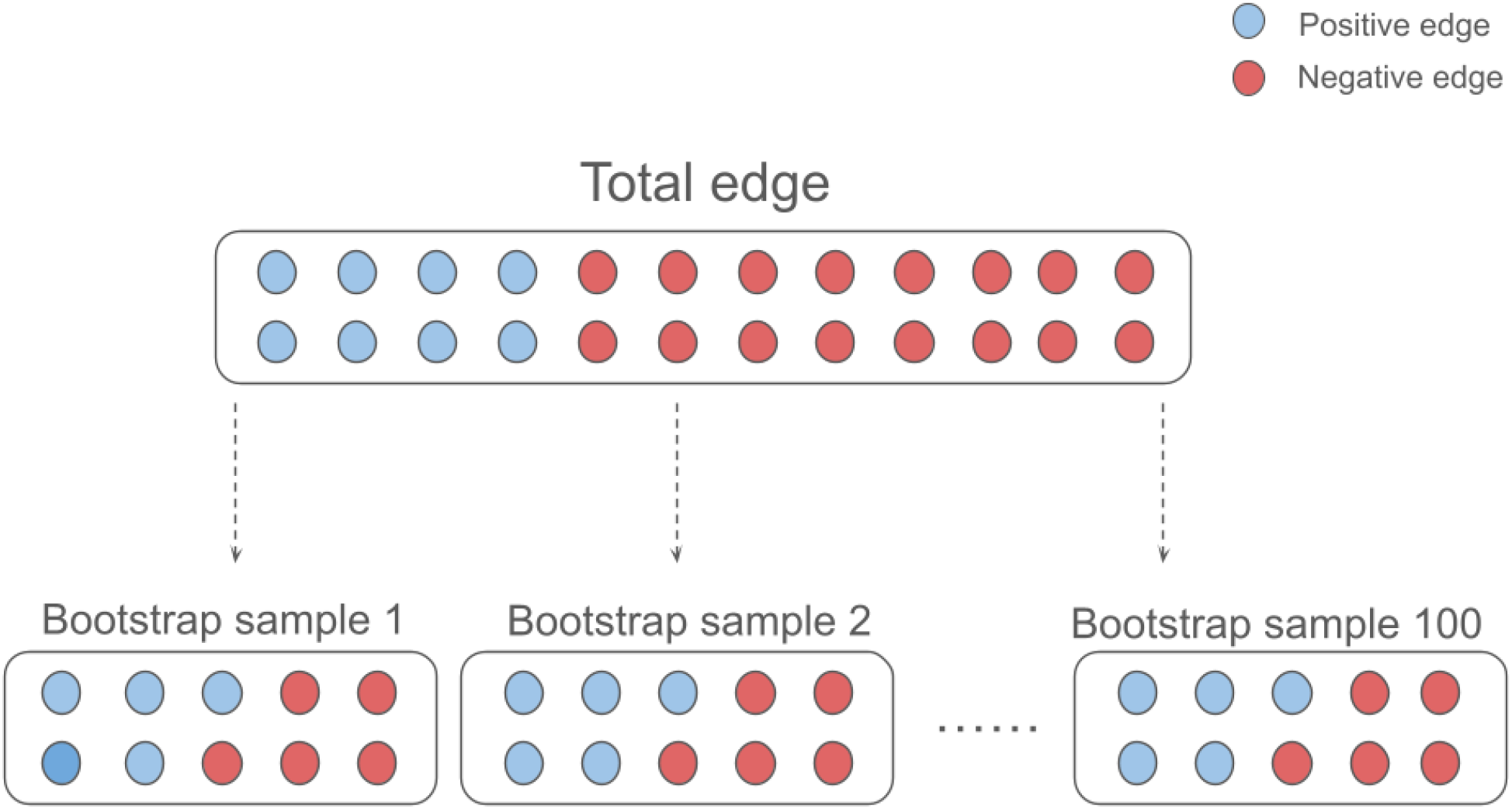
Bootstrap evaluation protocol used to obtain stable performance estimates. For each model–dataset–network combination, 100 bootstrap rounds are performed by retaining all positive test edges and randomly sampling an equal number of negative edges to form balanced test subsets. AUROC, AUPRC, and F1 scores are computed in each round, and the final reported metric is the average across all bootstrap iterations.

In each bootstrap iteration, the full set of positive test edges is retained, and an equally sized set of negative edges is randomly sampled without replacement to form a balanced evaluation subset. GRNPred computes area under the receiver operating characteristic curve (AUROC), area under the precision recall curve (AUPRC), and F1 score on this balanced bootstrap subset. The final reported metric for each dataset is the arithmetic mean across all 100 bootstrap rounds. This evaluation protocol mitigates bias induced by class imbalance, reduces metric variance across resamples, and enables confidence-consistent comparisons across network types and inference methods.

Under this evaluation protocol, GRNPred demonstrates strong and consistent predictive performance across all seven cell types and three regulatory network categories (Table 2). Test-set AUROC and AUPRC scores are consistently high, reflecting GRNPred’s ability to integrate multimodal biological features with transformer-based contextualization to accurately identify regulatory relationships. These results highlight the model’s robustness across varied ground truths and its adaptability to diverse expression landscapes.

**Table 1:**
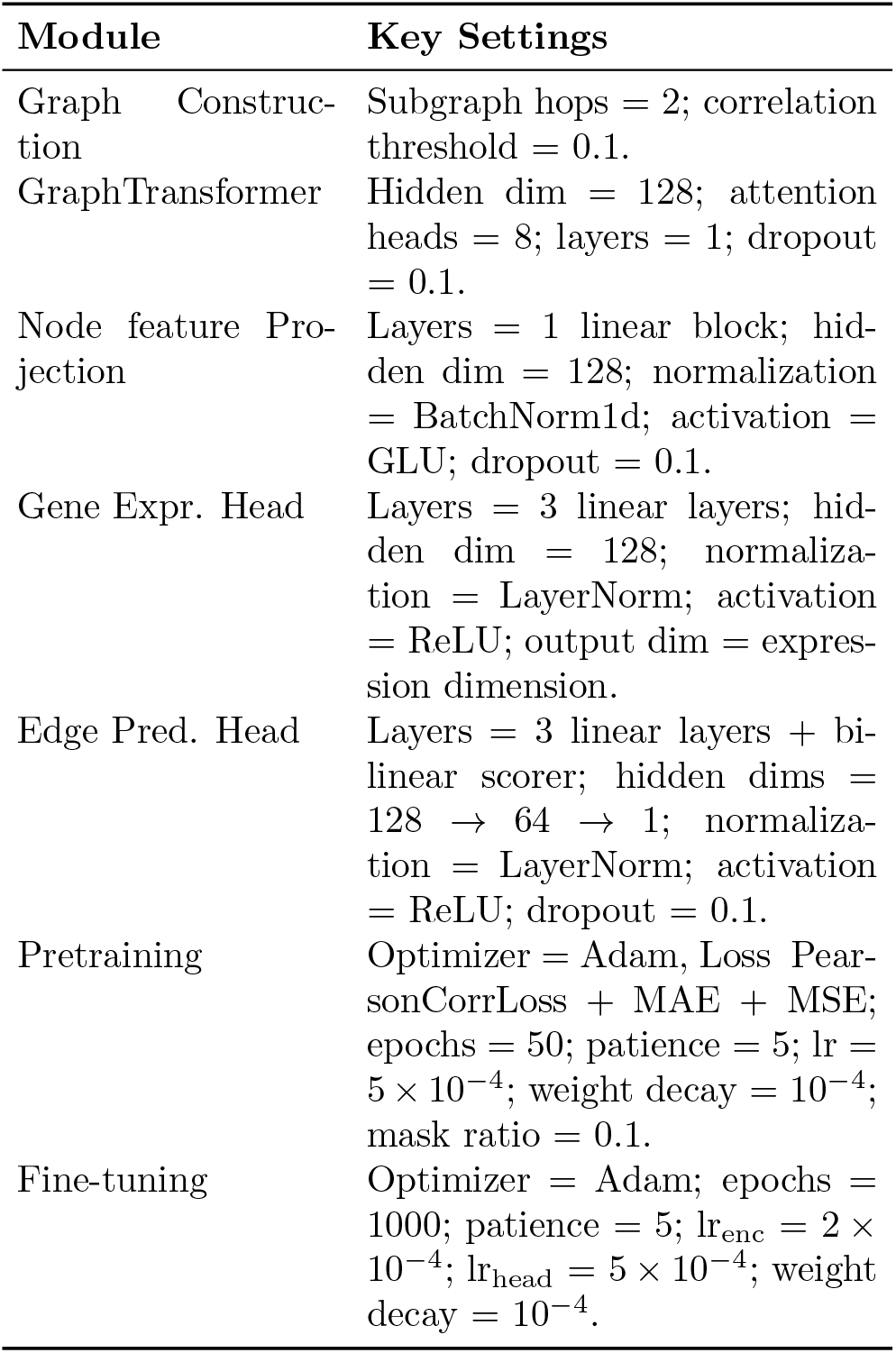
Hyperparameter configuration of GRNPred.

**Table 2:** Evaluation of performance of GRNPred across seven test datasets using three regulatory network constructions, reported as the average over tf500 and tf1000 TF–gene graph variants. Metrics include F1 score, AUROC, and AUPRC, assessed on nonspecific ChIP-seq, cell-type-specific ChIP-seq, and STRING GRNs.

| Dataset | Nonspecific ChIP-seq |  |  | STRING |  |  | Cell-type-specific ChIP-seq |  |  |
| --- | --- | --- | --- | --- | --- | --- | --- | --- | --- |
|  | F1 | AUROC | AUPRC | F1 | AUROC | AUPRC | F1 | AUROC | AUPRC |
| mHSC-GM | 0.92 | 0.94 | 0.93 | 0.94 | 0.97 | 0.96 | 0.96 | 0.97 | 0.96 |
| mHSC-l | 0.88 | 0.92 | 0.91 | 0.90 | 0.92 | 0.92 | 0.79 | 0.69 | 0.60 |
| mHSC-E | 0.83 | 0.83 | 0.79 | 0.80 | 0.86 | 0.87 | 0.86 | 0.77 | 0.71 |
| mESC | 0.84 | 0.84 | 0.88 | 0.88 | 0.89 | 0.92 | 0.93 | 0.92 | 0.84 |
| mDC | 0.87 | 0.92 | 0.92 | 0.87 | 0.93 | 0.93 | 0.88 | 0.89 | 0.88 |
| hESC | 0.84 | 0.88 | 0.87 | 0.88 | 0.94 | 0.94 | 0.80 | 0.77 | 0.83 |
| hHep | 0.85 | 0.91 | 0.89 | 0.85 | 0.92 | 0.91 | 0.84 | 0.80 | 0.74 |

Across the three types of GRNs, on STRING-based networks GRNPred achieves the highest inference accuracy, followed by non-specific ChIP-seq GRNs, with cell type-specific networks generally showing comparatively lower performance.

It is worth noting that the high accuracy of GRNPred is achieved under the setting that the numbers of positive and negative edges in the test dataset are equal.

### 2.3 GRNPred performs better than representative GRN inference methods

To contextualize the performance of GRNPred, we performed a comprehensive and carefully controlled comparative evaluation against five widely used and representative gene regulatory network (GRN) inference methods: GENELink [21], GNE (Gene Network Embedding) [22], STGRN (Spatial–Temporal Graph Regulatory Network) [23], GENIE3 [2], and GNNLink [17]. These methods provide a diverse and challenging benchmark set against which to assess the effectiveness, robustness, and generalization capability of GRNPred.

To ensure a strictly fair and reproducible comparison, all supervised competing methods were trained, validated, and evaluated using exactly the same edge splits as those employed for GRNPred. Ground-truth regulatory edges were stratified into 70% training, 15% validation, and 15% testing splits, with the proportion of positive and negative edges preserved across all subsets. Importantly, no model was exposed to test edges during training or hyperparameter tuning. Each baseline model was trained exclusively on the training split, with all hyperparameters selected solely on the basis of validation-set performance. Final evaluation metrics were computed on the held-out test split, ensuring that all reported results reflect true generalization rather than inadvertent information leakage or post hoc tuning.

Given the extreme class imbalance inherent to GRN inference—where true regulatory interactions constitute only a small fraction of all possible transcription factor–target pairs—all models were evaluated using the same standardized bootstrap procedure used to evaluate GRNPred. This ensures that GRNPred and all baseline methods are assessed under identical conditions and that the reported metrics are statistically stable and directly comparable.

Collectively, this evaluation framework ensures statistical rigor, methodological fairness, and direct comparability across models. By enforcing identical data splits, identical bootstrap testing, and consistent evaluation metrics, we provide an “apples-to-apples” comparison that isolates the contribution of each method’s modeling and representational capacity.

Across all evaluated network and cell types, GRNPred achieves the overall strongest performance in terms of both test-set AUROC and test-set AUPRC, demonstrating robust generalization across regulatory priors, and biological contexts (Figure 3). Under nonspecific ChIP-seq networks, which are known to be noisy and incomplete, GRNPred attains AUROC values ranging from approximately 0.83 to 0.94 and AUPRC values from approximately 0.79 to 0.93. In contrast, expression-driven methods such as GENIE3 and GNE generally exhibit lower performance and greater variability across datasets. This performance gap highlights GRNPred’s ability to effectively integrate weak or noisy regulatory signals to improve prediction accuracy.

**Figure 3:**
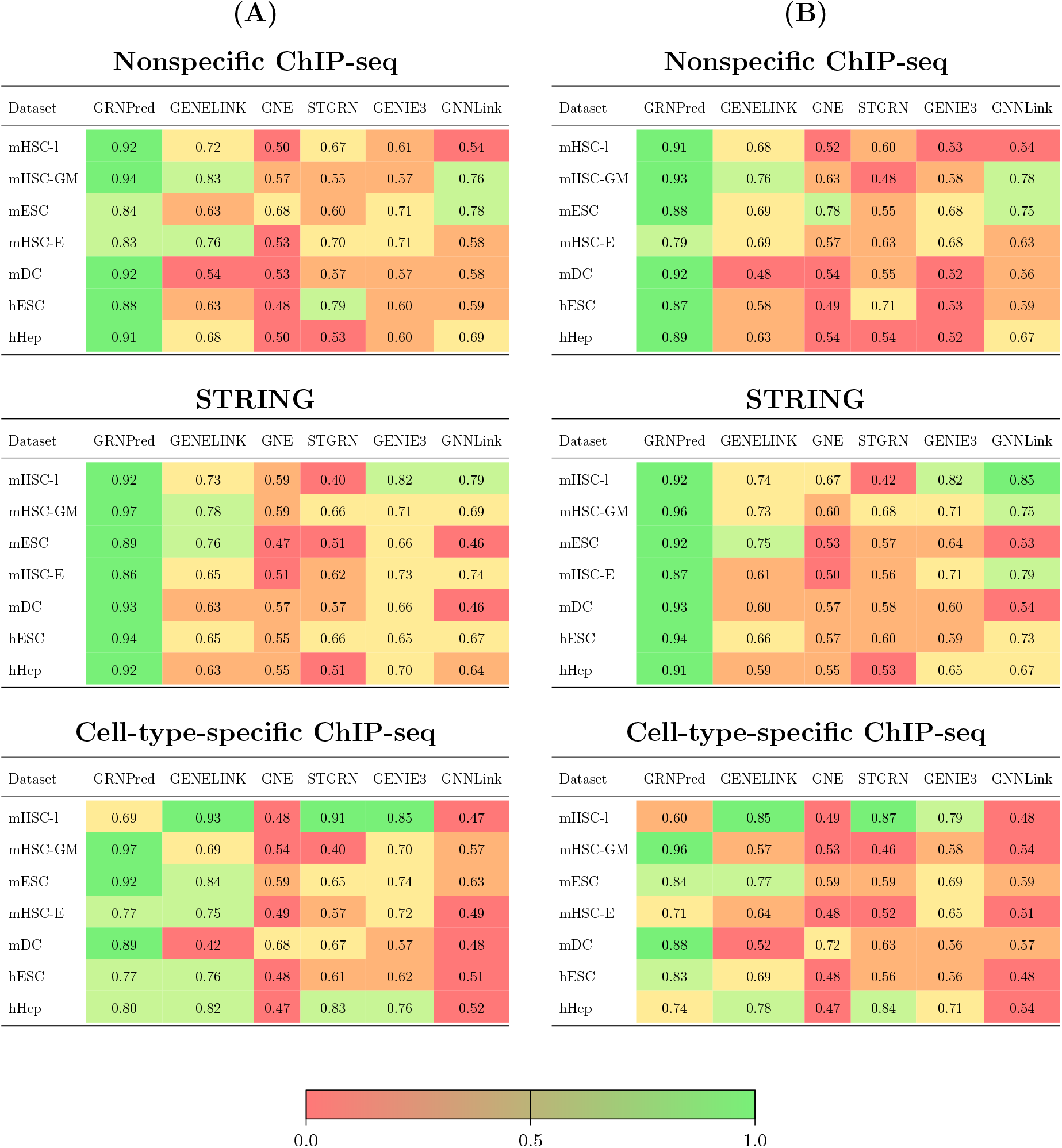
AUROC and AUPRC comparison of GRNPred against five representative GRN inference methods across seven datasets under three network constructions: nonspecific ChIP-seq, STRING, and cell-type-specific ChIP-seq. AUROC (A) and AUPRC (B) provide complementary assessments of ranking quality and precision under class imbalance. Results are averaged over the tf500 and tf100 variants of each dataset.

When evaluated using STRING-based network, overall performance improves for most approaches due to the higher relevance and specificity of the regulatory relationships. In this setting, GRN-Pred achieves competitive or superior performance on most datasets, with particularly strong results on mHSC-E, mESC, and hHep. While competing deep learning models—most notably STGRN and GNNLink—occasionally achieve strong performance on individual datasets, their results are often less stable across cell types. In contrast, GRNPred generally exhibits more consistent performance across datasets, indicating improved robustness to dataset-specific biases and regulatory heterogeneity.

Performance under cell-type–specific ChIP-seq networks constructions is notably more variable for all evaluated methods. In this setting, GRNPred demonstrates mixed performance across datasets. While it achieves strongest results on several datasets—such as mHSC-GM, mESC, mDC, and hESC—it performs less favorably on others, including mHSC-l, mHSC-E and hESC, where methods such as GENELink and STGRN attain higher AUROC and AUPRC values. Overall, GRNPred still performs better than all other methods on average.

Overall, these results demonstrate that GRN-Pred outperforms existing GRN inference methods in terms of overall robustness and consistency. The sustained gains in both AUROC and AUPRC across evaluation settings underscore the effectiveness of GRNPred’s multimodal design, which jointly integrates regulatory priors, global and local network topology, and learned representations to infer gene regulatory relationships in complex and heterogeneous biological systems.

### 2.4 Ablation study

To assess the contribution of each biological information source, we performed a unified modality ablation study in which individual node-level and edge-level feature types were independently removed from GRNPred. In each ablation, the corresponding modality was completely excluded from the model input, allowing us to quantify its standalone contribution to model performance. All ablations were evaluated on the tf500 non–cell-type-specific ChIP-seq network across seven cell types, using the same 70/15/15 data split and 100-round bootstrap testing procedure as the full model.

Table 3 reports the results of the unified modality ablation study evaluated using AUROC and AUPRC across seven datasets. The full GRNPred model demonstrates stable performance across diverse biological contexts, achieving baseline AUROC and AUPRC values averaging approximately 0.90 and 0.89, respectively. Removing individual feature modalities leads to performance changes—often degradation—in a dataset-dependent manner, indicating that distinct sources of biological information contribute complementary regulatory signals rather than serving as redundant inputs.

**Table 3:**
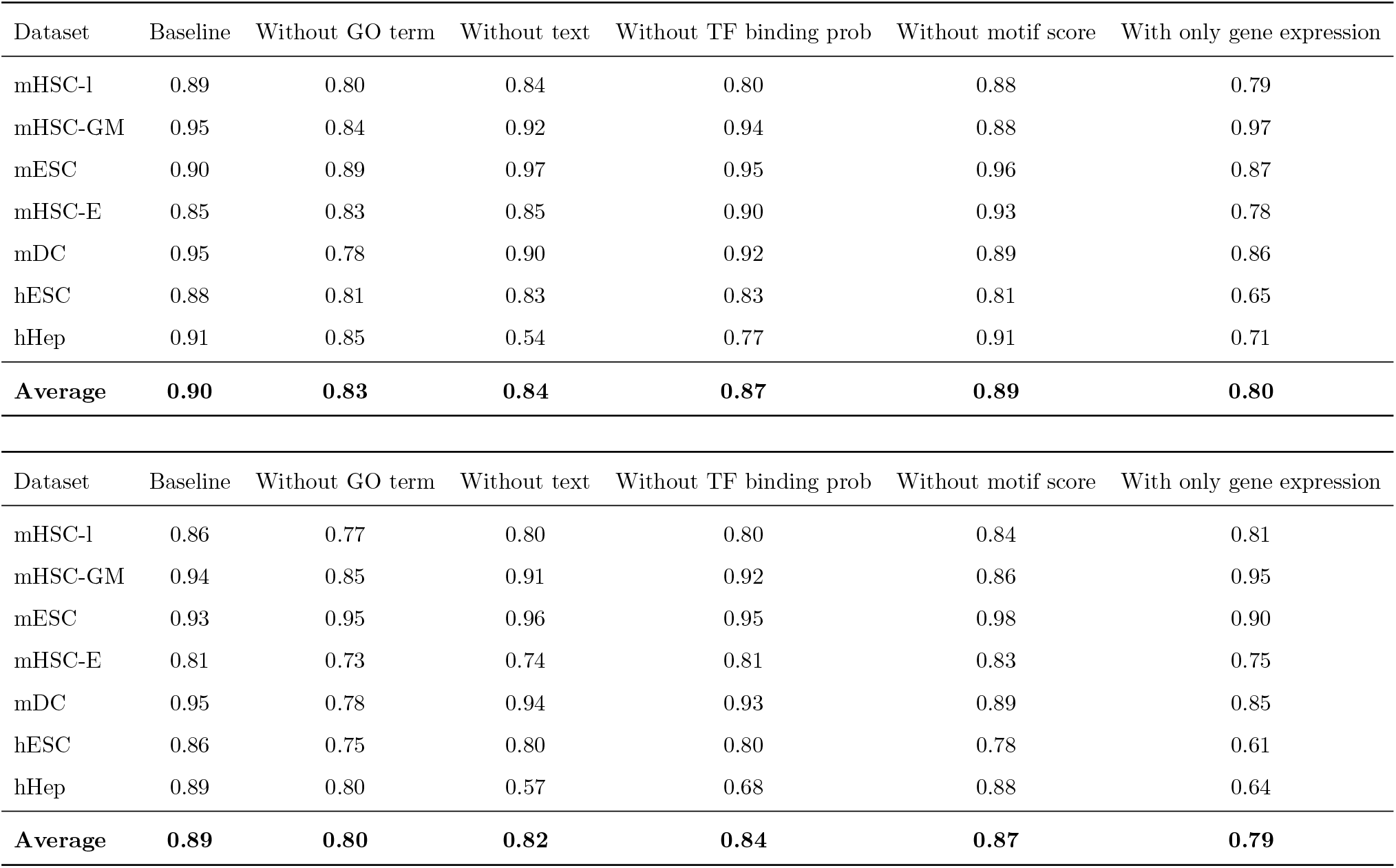
Ablation results (AUROC and AUPRC) across tf500 datasets of 7 cells.

In the extreme ablation setting where only gene expression is retained, overall performance is consistently lower than the full model across most datasets, with average AUROC and AUPRC values of 0.80 and 0.79, respectively. This result further supports the observation that expression-derived signals alone do not fully capture regulatory relationships, and that additional biological modalities provide complementary information. The extent of performance reduction varies across datasets, reflecting differences in the degree to which regulatory interactions can be inferred from expression patterns alone.

Among the evaluated modalities, Gene Ontology (GO) annotations exert a pronounced influence on model performance, particularly in hematopoietic datasets including mHSC-l, mHSC-GM, and mDC, where substantial reductions in both AUROC and AUPRC are observed following removal. In contrast, the effect of GO features is less prominent in embryonic stem cell datasets.

Text-based semantic features derived from curated gene descriptions exhibit highly heterogeneous effects across datasets. While their exclusion results in severe performance degradation in hHep—most notably a sharp decline in both AUROC and AUPRC—other datasets such as mESC show limited sensitivity or modest performance improvements. This variability indicates that textual information encodes context-specific regulatory cues that may be critical in certain biological systems but less informative in others.

Sequence-level priors, including transcription factor binding probability and motif scores, produce dataset-dependent effects that are often moderate but can be substantial in specific contexts. For example, removing TF binding probability reduces performance on mHSC-l and hHep, whereas removing motif scores yields large gains on mHSC-E.

This ablation analysis highlights the importance of multimodal integration for robust GRN inference. Functional annotations, semantic gene knowledge, and sequence-based priors each provide distinct and complementary regulatory evidence, and their relative contributions vary across biological contexts. These findings underscore the necessity of heterogeneous biological information fusion to achieve stable and generalizable regulatory network prediction.

### 2.5 Consistency between Ablation and SHAP Analyses

To further contextualize the ablation results, we conducted a group-level SHAP analysis [24] focusing on non-expression modalities (Table 4). Consistent with the ablation findings in Table 3, Gene Ontology (GO) features exhibit consistently high absolute SHAP values across most datasets, with particularly strong contributions in hematopoietic systems such as mHSC-l, mHSC-GM, and mDC. These results align with the pronounced reductions in AUROC and AUPRC observed when GO annotations are removed, confirming their role as a stable and informative source of functional regulatory constraints.

**Table 4:**
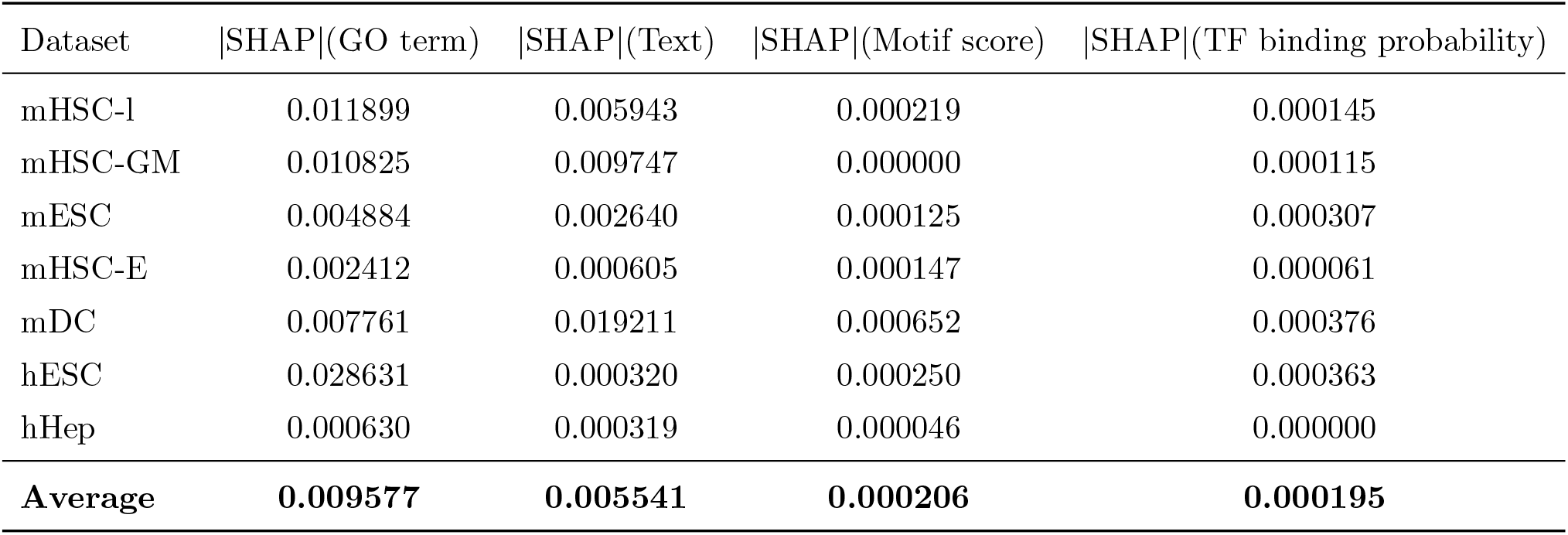
Group-level SHAP analysis of non-expression feature modalities across datasets. Absolute SHAP values indicate relative contribution magnitude.

Text-based semantic features show more heterogeneous SHAP contributions across cell types, closely mirroring their variable impact in the ablation study. In datasets such as mHSC-GM and hHep, text features exhibit relatively large absolute SHAP values, consistent with the substantial performance degradation observed upon their exclusion. In contrast, datasets such as mESC and mHSC-E display weaker SHAP contributions from text features, corresponding to limited sensitivity or modest performance changes in the ablation setting. These patterns suggest that textual information captures context-dependent regulatory cues that are critical in certain biological systems but less informative in others.

In comparison, sequence-level priors, including motif scores and TF binding probabilities, consistently show low absolute SHAP magnitudes across datasets. This indicates that while these features may refine regulatory confidence in specific contexts, they play a secondary role relative to functional and semantic information. Overall, the SHAP analysis corroborates the ablation results by demonstrating that GRNPred leverages complementary, non-redundant biological modalities, with their relative importance varying across cellular contexts rather than being dominated by any single information source.

### 2.6 Biological Interpretation of Predicted Gene Regulatory Networks

To assess the biological relevance of GRNPred predictions, we analyzed transcription factor–target gene pairs with predicted probabilities greater than 0.5, which were considered high-confidence regulatory interactions. For the tf 500 mHSC-GM non-specific ChIP-seq dataset, this filtering produced a compact regulatory subnetwork enriched for genes involved in cell-cycle progression, DNA replication, metabolic regulation, and immune signaling (Figure 4).

**Figure 4:**
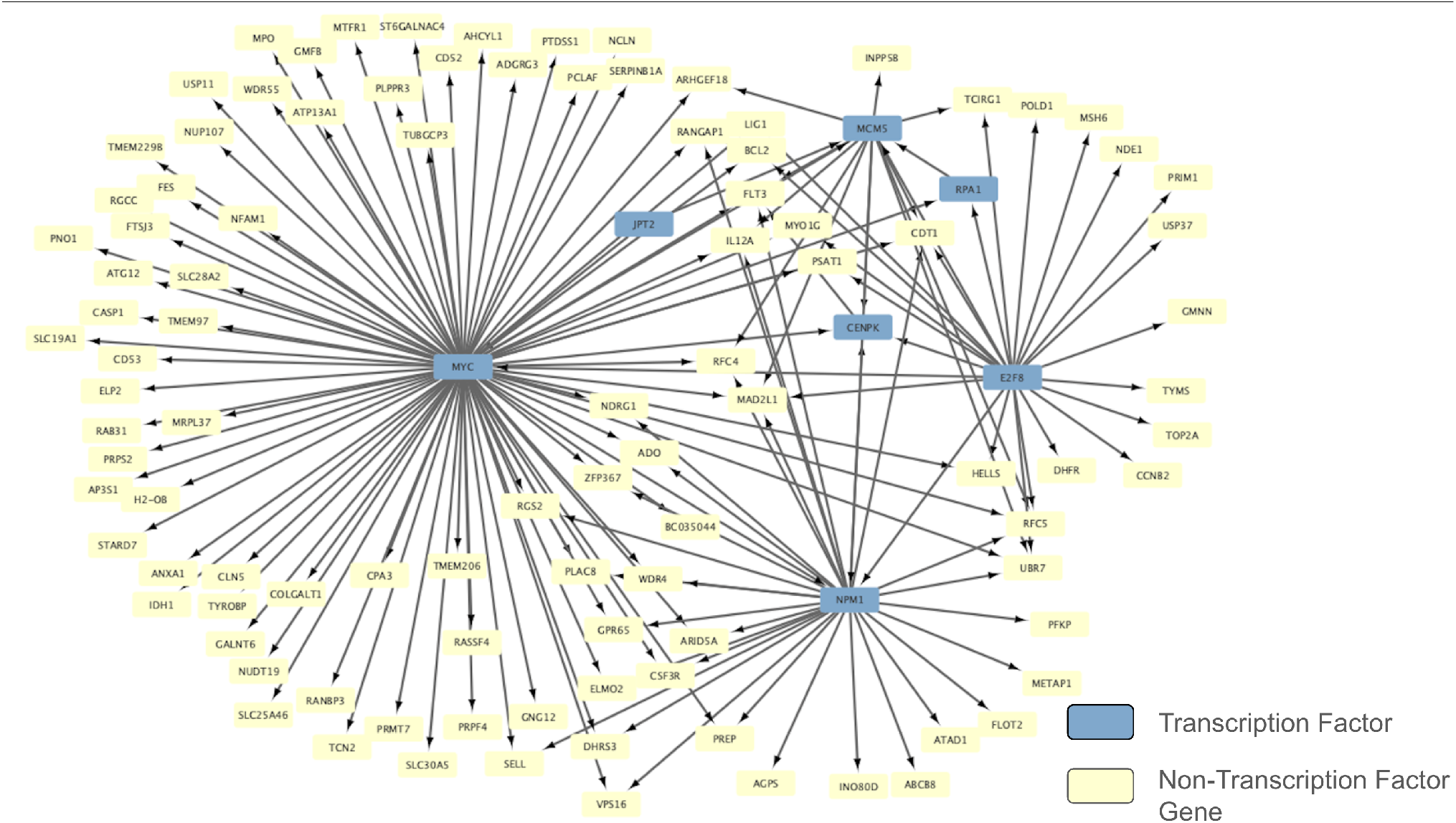
High-confidence gene regulatory subnetwork inferred by GRNPred in mHSC-GM cells for the tf 500 non-specific ChIP-seq dataset. The network was constructed by retaining transcription factor–target gene interactions with predicted probabilities greater than 0.5.

Within this network, *MYC* emerged as a dominant regulatory hub. *MYC* is a well-established master regulator of cellular growth and proliferation and plays a central role in controlling DNA replication, metabolism, and cell-cycle progression in hematopoietic progenitor cells [25, 26]. Consistent with these known functions, GRNPred predicted high-confidence interactions between *MYC* and multiple replication- and cell-cycle–associated genes, including *MCM5, RPA1, LIG1, MAD2L1*, and *CENPK*, which are consistent with their essential roles in DNA synthesis and chromosomal stability [27, 28, 29, 30]. In addition, *MYC* was predicted to regulate metabolic and immune-related genes such as *PRPS2, PSAT1, FLT3*, and *CSF3R*, supporting its reported role in coordinating proliferative and signaling programs during hematopoietic development [31, 32].

The predicted network also highlighted cell-cycle regulatory modules involving *E2F8*, a member of the E2F transcription factor family known to control S-phase entry and DNA replication [33]. Predicted *E2F8* targets included genes such as *TOP2A, TYMS, POLD1*, and *MAD2L1*, consistent with their established roles in DNA replication, nucleotide biosynthesis, and mitotic check-point control [34, 35, 36, 29].

## 3 Discussion

In this study, we introduce GRNPred, a multimodal graph transformer framework for gene regulatory network inference that combines heterogeneous biological evidence with a two-stage training paradigm. GRNPred first undergoes self-supervised pretraining, during which the model learns to reconstruct masked gene-expression profiles within TF-centered subgraphs, drawing inspiration from masked autoencoding approaches such as BERT [12] and graph-level self-supervised methods like GraphMAE [13]. It then goes through supervised fine-tuning on regulatory edge prediction, allowing the pretrained encoder to specialize its learned representations toward identifying TF–gene interactions. Our experiments demonstrate that this pretrain–finetune strategy substantially improves predictive accuracy and stability, consistent with findings in representation learning for graphs [5, 4, 9].

Across seven human cell types and three distinct ground-truth regulatory networks, GRNPred consistently outperformed representative graph-based and machine learning methods. GRNPred’s strong performance indicates that the representations learned during self-supervised expression modeling transfer effectively to downstream regulatory prediction tasks, consistent with broader observations in graph representation learning [10]. A central insight from our results is the importance of multimodal biological integration within this pretraining–finetuning framework. The ablation study revealed that each feature class—gene expression, GO annotations [15, 37], semantic gene embeddings derived from UniProt text [38], motif scores computed from JASPAR PWMs [39], and TF-binding probabilities—provides unique and complementary information.

Nevertheless, GRNPred has several limitations. The model relies heavily on the availability of high-quality biological annotations, including curated gene text descriptions and accurate GO-term mappings. Incomplete, inconsistent, or missing annotations can diminish the utility of semantic and functional modalities. Additionally, the computation of certain edge-level features—such as motif scores and TF-binding probability estimates—requires substantial preprocessing time and external tools such as FIMO [40] and JASPAR [39], which may restrict scalability in environments lacking these resources. The current pretraining objective focuses exclusively on expression reconstruction; incorporating multi-task or contrastive objectives may further strengthen the model’s ability to learn intrinsic regulatory structure, as suggested in recent self-supervised GNN research [13, 9].

A more fundamental limitation lies in the fact that GRNPred is trained independently on each dataset. While this prevents information leakage, it also means the model does not exploit shared regulatory principles across related cell types or conditions. As a consequence, GRNPred exhibits limited cross–cell type generalization. Extending the framework to support multi-task learning, transfer learning, or shared-encoder architectures could allow the model to capture common regulatory logic while preserving cell-type-specific distinctions.

Looking ahead, GRNPred’s combination of self-supervised transcriptional pretraining and supervised regulatory finetuning—implemented within a multimodal graph-transformer architecture [9, 10]—provides a strong foundation for further methodological development. Future extensions may explore richer self-supervised objectives, improved multimodal fusion mechanisms, or adaptive subgraph sampling strategies to better capture long-range dependencies in regulatory circuits. Incorporating scalable training across multiple datasets, rather than treating each cell type independently, could enable the model to learn shared regulatory principles and improve cross– cell-type generalization. Additionally, integrating more diverse biological modalities, automating feature extraction pipelines, and reducing reliance on time-intensive preprocessing steps (such as motif scoring via FIMO [40]) may further enhance the utility and practicality of GRNPred. As these directions progress, GRNPred offers a promising platform upon which more advanced and biologically expressive GRN inference frameworks can be built.

## 4 Methods

### 4.1 Creating gene co-expression networks

To generate a gene co-expression graph used as part of the *GRNPred* input, we first computed pairwise Pearson correlation coefficients between the expression values of any two genes across all gene–gene expression profiles within each dataset. Pearson correlation is widely used in transcriptomic analysis and gene regulatory network construction due to its simplicity, interpretability, and effectiveness in capturing linear co-expression relationships arising from shared regulatory control and coordinated transcriptional programs [41, 42]. Specifically, for the expression profiles (*g*_*i*_, *g*_*j*_) of two genes *i* and *j*, the Pearson correlation coefficient is defined as

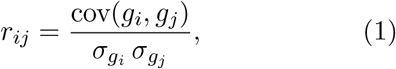

where cov(·, ·) denotes the covariance between expression profiles and *σ*_*g*_ represents the standard deviation of a gene’s expression across samples. This normalization accounts for gene-specific variation in expression magnitude and enables consistent comparison across heterogeneous expression distributions [43].

Following computation of the full correlation matrix, we applied a thresholding step to construct a sparse and biologically meaningful co-expression network (Figure 5). All gene pairs with correlation values below 0.1 were discarded to suppress weak or noisy associations that are unlikely to reflect true transcriptional coordination. Correlation-based thresholding is commonly adopted in GRN inference to balance sensitivity and specificity and to prevent dense, noise-dominated graphs that degrade downstream modeling performance [44].

**Figure 5:**
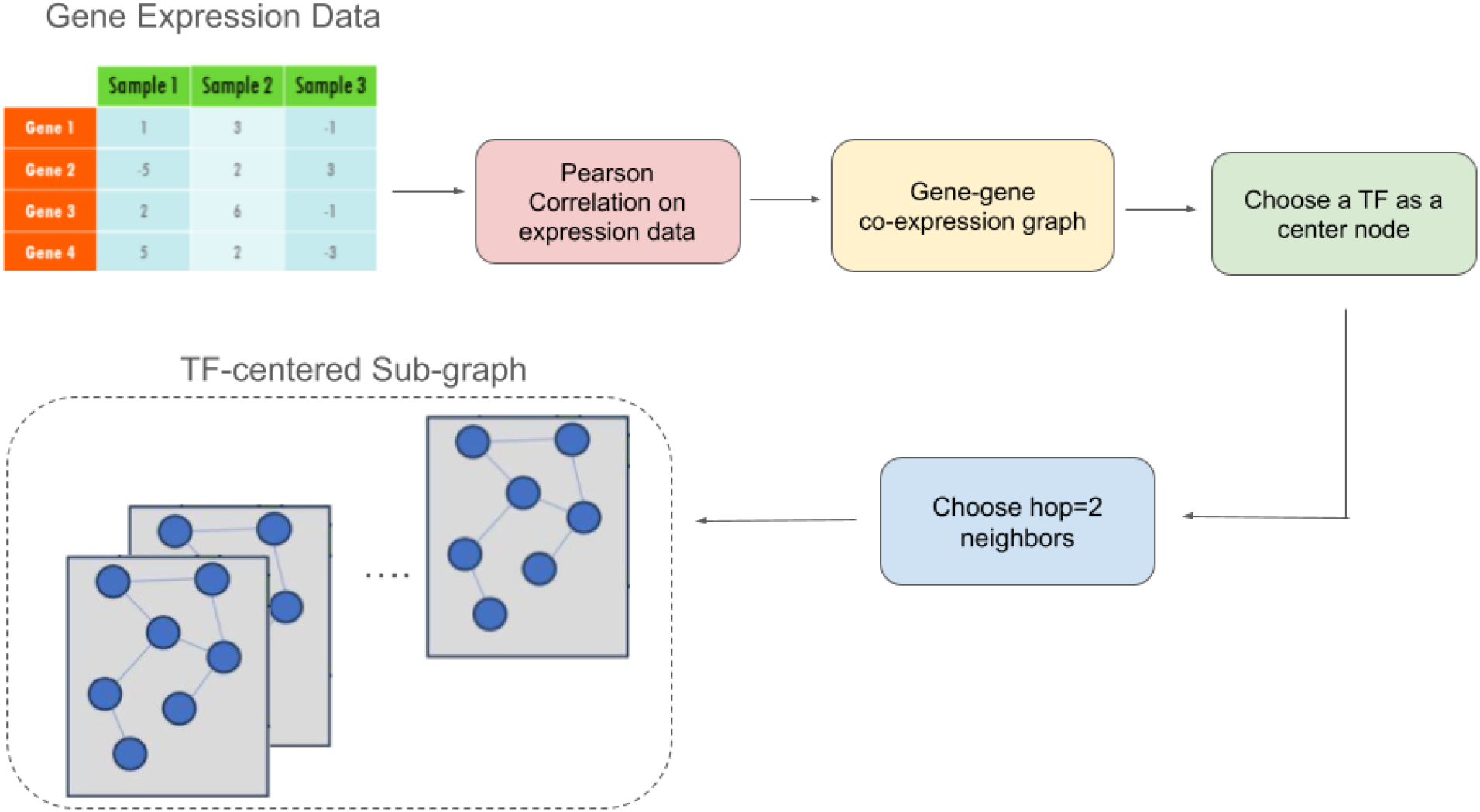
Subgraph construction pipeline for GRNPred. A global co-expression graph is first built using pairwise Pearson correlation, with weak associations removed by thresholding. TF-centered 2-hop subgraphs are then extracted from the resulting sparse graph and used as model inputs.

### 4.2 Multimodal MLP Encoder

Each TF-centered subgraph contains nodes corresponding to genes enriched with multiple heterogeneous feature channels, including expression measurements, Gene Ontology (GO) functional description, LLaMA-based text embeddings derived from curated protein annotations, and sequence-level TF-binding features (see Section Dataset and Input Features for details). These modalities capture complementary biological evidence spanning transcriptional activity, functional semantics, and regulatory potential. However, they differ substantially in scale, dimensionality, sparsity, and semantic interpretation, which makes naïve feature concatenation suboptimal for downstream graph-based learning.

To effectively integrate these heterogeneous inputs, GRNPred employs a multimodal MLP encoder that explicitly aligns and convert raw feature channels into a unified latent representation prior to message passing. For each gene *i*, the modality-specific feature vectors are concatenated to form an initial representation 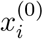. This vector is first batch-normalized to reduce scale imbalance across modalities and to improve optimization stability during training.

The normalized representation is then projected using a gated linear unit (GLU), which enables adaptive, modality-dependent feature selection:

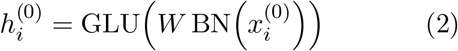

The GLU introduces learnable gates that modulate feature activations, allowing the encoder to suppress noisy or redundant signals while emphasizing biologically informative dimensions. This gating mechanism is especially important in multimodal biological settings, where certain modalities may be informative only for specific TFs, cell types, or regulatory contexts. Compared to standard nonlinear activations, the GLU provides a more expressive fusion strategy by decoupling feature transformation from feature selection [45].

The resulting embeddings 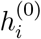 reside in a shared space with consistent scale and dimensionality, making them suitable as initial node representations for transformer-based message passing.

An analogous MLP encoder is applied to raw edge features, including motif enrichment scores and TF-binding probabilities, to generate initial edge embeddings. Encoding edge attributes separately allows regulatory evidence to be incorporated directly into attention computations without conflating node- and edge-level information.

### 4.3 GraphTransformer: Attention-Based Regulatory Reasoning

At the core of GRNPred is a multi-layer Graph-Transformer built using TransformerConv layers [46] (Figure 1), which extend transformer-style self-attention mechanisms to graph-structured data. This architecture enables the model to perform context-aware regulatory reasoning by dynamically modulating information flow between genes based on node-level representations. In contrast to conventional message-passing graph neural networks (GNNs), which aggregate information using fixed or uniformly parameterized neighborhood operators, the GraphTransformer allows each node to selectively attend to its neighbors in a data-dependent manner.

This adaptive attention mechanism is particularly well suited for gene regulatory network inference, where the strength, relevance, and directionality of regulatory interactions vary substantially across transcription factors, target genes, and cellular contexts. By assigning higher attention weights to biologically informative neighbors, the GraphTransformer can prioritize real regulatory signals while suppressing spurious associations.

At layer *t*, node representations are updated by attending over their local neighborhoods:

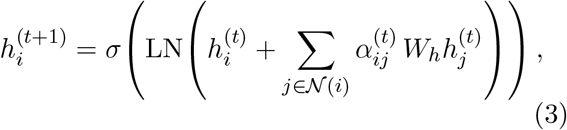

where 𝒩 (*i*) denotes the set of neighboring nodes of node *i*, 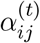 are attention coefficients, and *W*_*h*_ is a trainable linear projection matrix. The attention coefficients are computed using an edge-aware attention mechanism:

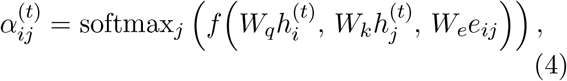

where *W*_*q*_, *W*_*k*_, and *W*_*e*_ are learnable projection matrices, *e*_*ij*_ denotes the edge embedding associated with the interaction between nodes *i* and *j*, and *f* (·) is a compatibility function implemented as scaled dot-product attention. By incorporating edge embeddings into the attention computation, regulatory priors such as motif enrichment scores and transcription factor binding probabilities directly modulate message propagation. This edge-aware attention mechanism enables GRNPred to capture regulatory directionality and interaction strength within the attention process itself, rather than relying solely on post-hoc feature fusion.

Residual connections and layer normalization (LN) are employed to stabilize optimization and facilitate the training of deeper transformer-based architectures. Specifically, layer normalization is applied to normalize node representations within each layer, ensuring that feature statistics remain well-conditioned across layers and preventing the accumulation of representation drift during message passing. In combination with residual connections, this design mitigates gradient degradation and enables stable information propagation through multiple GraphTransformer layers. The nonlinear activation function *σ*(·) is implemented using the rectified linear unit (ReLU), which introduces nonlinearity into the learned representations and enables the model to capture complex regulatory dependencies present in transcriptional networks.The importance of nonlinear activation functions for learning high-capacity representations in deep neural networks has been extensively established in prior work [47, 48].

### 4.4 MLP Decoder Head for Gene Expression

We employ a node-wise multilayer perceptron (MLP) as the gene expression decoder. Given the final node representation 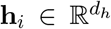 produced by the graph encoder, the decoder maps it to a reconstructed gene expression vector 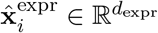:

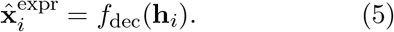

The decoder consists of a Layer Normalization followed by two hidden linear layers with ReLU activations and a final linear projection:

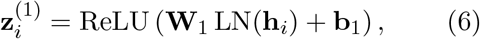

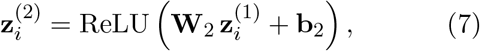

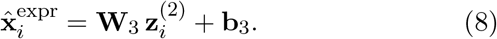

Here, LN(·) denotes Layer Normalization, and {**W**_*k*_, **b**_*k*_} are learnable parameters. The final output dimension *d*_expr_ matches the dimensionality of the gene expression features, enabling full-vector prediction for each node.

This decoder operates independently on each node embedding, providing a direct mapping from the learned latent space to the original gene expression space.

### 4.5 MLP Decoder for TF–Gene Regulatory Prediction

Following message passing through the Graph-Transformer encoder, GRNPred evaluates transcription factor–gene regulatory potential using an MLP-based edge decoder. For each candidate regulatory pair (*i, j*), corresponding to a transcription factor and a potential target gene, the decoder constructs a composite feature representation that captures multiple complementary perspectives of their interaction. Specifically, this representation concatenates the node embeddings (*h*_*i*_, *h*_*j*_), their element-wise product *h*_*i*_ ⊙ *h*_*j*_, and their absolute difference |*h*_*i*_−*h*_*j*_|. This feature construction strategy is widely adopted in representation learning for relational and link prediction tasks, as it jointly encodes similarity, interaction strength, and directional asymmetry [5, 22].

The inclusion of both multiplicative and difference-based terms allows the decoder to capture regulatory patterns that cannot be represented by simple linear similarity. The element-wise product emphasizes shared activity patterns and co-regulated signals between transcription factors and target genes, while the absolute difference captures feature disparities that may indicate directional regulatory effects. By combining these complementary representations, the model can capture asymmetric similarity patterns relationships, which has been shown to improve performance in graph-based prediction tasks [49].

The final regulatory score is computed as

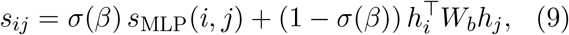

where *s*_MLP_(*i, j*) denotes the output of the nonlinear MLP decoder applied to the composite edge representation, *W*_*b*_ is a learnable bilinear weight matrix, and *β* is a trainable scalar that balances the two terms. Bilinear scoring functions are commonly used in relational learning as a structured similarity prior that captures global compatibility between paired embeddings [50], while MLP-based decoders enable nonlinear integration of heterogeneous features and contextual information [17].

The learnable mixing coefficient *σ*(*β*) allows the model to balance between the nonlinear MLP score and the bilinear interaction score. This adaptive weighting enables the decoder to combine expressive feature integration with structured similarity between node embeddings. Such a design is useful for gene regulatory network inference, where regulatory interactions can arise from both direct transcription factor binding and more complex context-dependent regulatory effects. By combining these two complementary signals, the decoder improves its ability to distinguish true regulatory relationships from spurious associations.

### 4.6 Dataset and Input Features

All experiments are conducted on the BEELINE benchmarking suite of gene regulatory network (GRN) datasets, which provides curated single-cell expression matrices, transcription factor (TF) lists, and three categories of ground-truth regulatory networks (non–cell-type-specific ChIP-seq, cell-type-specific ChIP-seq, and STRING-based interactions). The benchmark includes seven cell types across human and mouse (hESC, hHep, mDC, mHSC-GM, mHSC-E, mHSC-L, and mESC). For each cell type, we construct two graph variants by combining the curated TF list with the top 500 or top 1000 highly variable genes, following the BEELINE [19] preprocessing protocol; here, highly variable genes are defined as those exhibiting the largest expression variance across single cells and are typically enriched for regulatory activity and informative transcriptional dynamics. GRNPred operates directly on the raw expression values provided by BEELINE without additional normalization, preserving the original signal structure.

To construct an initial co-expression scaffold, we compute pairwise Pearson correlation coefficients for all gene pairs within each dataset. Edges with correlation magnitude below 0.1 are removed to suppress weak or noisy associations. The resulting sparse graph is then used to extract TF-centered 2-hop subgraphs that serve as inputs to GRNPred.

In addition to gene expression values and co-expression network structure, GRNPred incorporates multiple external biological resources to construct multimodal node and edge features. Gene-level function description text is obtained from the UniProt Knowledgebase, a curated repository of protein sequence and functional annotation [38]. These textual descriptions are embedded using a LLaMA-based encoder [16] to provide semantic context complementary to expression-derived features.

Moreover, sequence-based regulatory features are derived from the human reference genome and gene annotations hosted at NCBI as follows. Promoter sequences of the genes are extracted from the human reference assembly provided by the Genome Reference Consortium and distributed via NCBI [51]. Gene coordinates and transcription start sites (TSSs) are defined using the GENCODE comprehensive gene annotation set [52]. For each gene, we define the promoter region as the 200 bp sequence immediately upstream of its TSS. TF-binding motif scores are computed by scanning these promoter sequences using FIMO, the motif-scanning tool from the MEME Suite that applies a log-likelihood–based statistical framework to Position Weight Matrix (PWM) search [40], against the binding motifs (i.e., PWMs) of the TFs obtained from the JASPAR 2024 database [39]. For each TF–gene pair, the highest FIMO-reported motif match score within the promoter window is retained as the motif-based edge feature. From the same 200 bp window, we additionally compute a TF-binding probability estimate as an feature, using a PWM-based occupancy model that integrates motif match strength, local nucleotide composition, and background base frequencies estimated from the GRCh38 assembly [51].

Furthermore, functional biological priors are incorporated through Gene Ontology (GO) annotations. The GO structure and term definitions are obtained from the official OBO-format ontology released by the Gene Ontology Consortium [37]. The OBO file is parsed using the obonet library to map GO terms to genes; these term sets are encoded as multi-hot vectors and projected into dense embeddings before being integrated into the multimodal node representation. For genes without any mapped GO terms, the corresponding GO feature vector is set to zero, ensuring a consistent feature dimensionality across all nodes. GO features provide functional context that complements expression- and sequence-based signals during regulatory inference.

Taken together, the BEELINE expression datasets enriched with UniProt text descriptions [38], NCBI-derived genomic resources [51, 52], JASPAR PWMs for motif scoring [39], and Gene Ontology annotations [37] form a multimodal feature representation on which GRNPred operates. This combination of expression, sequence, functional, and semantic information provides a biologically grounded foundation for transformer-based GRN reconstruction.

### 4.7 Training

GRNPred is trained using a two-stage protocol that combines self-supervised representation learning with supervised regulatory inference. This design separates representation learning from task-specific supervision, allowing the model to first capture general biological structure before specializing in TF–target interaction prediction. Specifically, the two stages of the training process are defined as (i) self-supervised pretraining via masked gene-expression reconstruction and (ii) supervised finetuning for TF–target edge classification (Figure 1).

Both training stages operate on TF-centered two-hop subgraphs extracted from the underlying co-expression. All experiments adhere to the fixed 70%/15%/15% train, validation, and test edge splits described above to ensure a strictly controlled evaluation protocol. Models are trained independently for each dataset and for each graph variant (tf500 and tf1000), with no parameter sharing across datasets, cell types, or experimental conditions. This setup avoids information leakage and allows performance comparisons to reflect dataset-specific regulatory characteristics.

#### 4.7.1 Stage I: Self-Supervised Pretraining to Predict Masked Gene Expression Values

The objective of the pretraining stage is to learn robust node representations that capture local regulatory context and multimodal biological structure without relying on labeled TF–target interactions. During this stage, the model is optimized to reconstruct masked gene-expression values within each TF-centered subgraph input, following the principles of masked autoencoding widely used in self-supervised learning [12, 53, 13]. This formulation encourages the GraphTransformer to encode informative dependencies among neighboring genes, building on recent advances in transformer-based graph representation learning.

For each training iteration, the training process iterates over TF-specific subgraphs and constructs a subgraph-level `Data` object containing node features, edge indices and edge attributes.

A fixed fraction of nodes within each TF-centered gene co-expression subgraph is randomly selected for masking. For both training and validation, a mask ratio of 0.1 is used. For masked nodes, the raw gene expression feature channels are set to zero, while all other feature modalities remain unchanged. The model is then trained to reconstruct the original gene expression vector of each masked node using contextual information propagated through the GraphTransformer. In other words, the model predicts the full expression profile of the masked gene based on information from neighboring genes within the co-expression subgraph and the available multimodal biological features. To prevent trivial solutions and ensure that predictions rely on meaningful network context, the reconstruction loss is computed only for nodes that are both (i) masked and (ii) connected to at least one edge within the gene co-expression subgraph.

The reconstruction objective is defined as an unweighted combination of three terms: mean squared error (MSE), mean absolute error (MAE), and a Pearson correlation–based loss. Each component contributes equally to the overall objective, balancing sensitivity to absolute magnitude (MSE), robustness to outliers (MAE), and preservation of relative expression patterns (Pearson correlation). In particular, the Pearson correlation–based loss measures the linear agreement between the predicted and target expression profiles after centering and variance normalization, making it invariant to global scale and offset differences. As a result, this term explicitly encourages the model to recover correct co-variation and ordering of expression values, even when absolute amplitudes differ. The composite loss formulation avoids overfitting to any single error metric and promotes stable learning of expression-aware representations that generalize across datasets with heterogeneous expression scales and noise characteristics.

Validation during pretraining follows the same masking strategy and is performed on held-out validation subgraphs and edges. Early stopping [54] is applied based on the minimum validation reconstruction loss, and the checkpoint with the best validation performance is retained for subsequent finetuning.

#### 4.7.2 Stage II: Supervised Finetuning for GRN prediction

Following self-supervised pretraining, GRNPred is finetuned in a supervised manner for TF–gene regulatory edge prediction. At this stage, the learning objective shifts from masked expression reconstruction to discriminative modeling of regulatory interactions, enabling the model to directly separate true regulatory edges from non-regulatory gene pairs. The pretrained Graph-Transformer encoder is used as initialization, allowing GRNPred to leverage contextual representations learned from unlabeled data during pretraining while efficiently adapting to the downstream supervised task.

During finetuning, the same TF-centered two-hop subgraphs are used as input to ensure architectural and representational consistency across training phases. In contrast to the self-supervised stage, node masking is entirely disabled, and the mask ratio is fixed to zero for both training and evaluation. As a result, the encoder operates on fully observed multimodal node features, including gene expression profiles and external biological annotations. In addition, each node is augmented with a learnable positional (node identity) embedding, which is concatenated with the original multimodal features before being passed to the encoder. This positional encoding provides a consistent node-level reference across subgraphs, allowing the model to distinguish individual genes and maintain structural awareness beyond local message passing. Finetuning therefore focuses on relational reasoning under complete biological context, while the decoder learns to map contextualized node embeddings to regulatory interaction scores.

The GraphTransformer encoder and the MLP-based edge decoder are optimized jointly using a single AdamW optimizer configured with two distinct parameter groups, allowing separate learning dynamics for pretrained components and task-specific classification layers. Encoder parameters, including multimodal projection layers, positional encodings, stacked `TransformerConv` layers, are trained with a conservative base learning rate of 2 × 10^−4^ to preserve generalizable representations learned during pretraining. In contrast, the edge-classification head is trained with a higher learning rate of 5 × 10^−4^, enabling faster convergence under supervised objectives without destabilizing the encoder. GRNPred is trained using a combination of a pairwise AUC loss and an average precision (AP) surrogate loss, both computed on the predicted edge logits. The AUC loss encourages correct ranking between positive and negative edges, while the AP loss improves precision–recall performance under severe class imbalance. The final objective is a weighted combination of these two losses, aligning optimization with evaluation metrics used in GRN inference.

To further stabilize optimization and mitigate overfitting, learning rates are dynamically adjusted using a `ReduceLROnPlateau` [55] scheduler monitored on validation performance. When validation metrics fail to improve within a predefined patience window, the scheduler reduces the learning rate to allow finer-grained optimization in later training stages. This adaptive scheduling strategy is particularly important given the severe class imbalance and dataset heterogeneity characteristic of GRN inference tasks.

Validation is performed on held-out regulatory edges from the validation split. Model selection is based exclusively on validation-set performance, and the best-performing checkpoint is retained for final evaluation. The selected model is then evaluated on the held-out test edges using the same bootstrap-based testing protocol.

#### 4.7.3 Optimization Details

Both training stages employ the AdamW optimizer with a weight decay of 1 × 10^−4^ and a `ReduceLROnPlateau` scheduler configured to reduce the learning rate by a factor of 0.5 when the validation loss stagnates for two epochs, with a minimum learning rate of 1 × 10^−6^. Mixed-precision training is enabled using automatic mixed precision (AMP) together with gradient scaling to improve computational efficiency and numerical stability on GPU hardware. During edge-level finetuning, gradient norms are clipped to a maximum value of 2.0 to mitigate optimization instability in the deep transformer-based architecture.

Across both pretraining and finetuning stages, early stopping is applied using a fixed patience window, and model parameters are restored from the checkpoint achieving the best validation performance. This two-stage training protocol—self-supervised expression reconstruction followed by supervised edge-level finetuning—produces a GRNPred model that is simultaneously grounded in transcriptional structure and highly discriminative for regulatory interaction prediction.

### 4.8 Benchmarking Setup

We conducted a structured benchmarking study to evaluate GRNPred against a curated set of established and recent gene regulatory network inference approaches spanning classical statistical models, graph-based methods, and deep learning architectures. The primary goal of this evaluation was to characterize relative performance under controlled and reproducible conditions, while eliminating confounding effects introduced by inconsistent preprocessing, data partitioning, or evaluation procedures.

All the GRN inference models were run using their publicly released reference implementations, adhering to parameter choices and training protocols recommended in their original publications. Where applicable, documented preprocessing pipelines were followed exactly as described in the original implementations, and all methods were applied to the same input datasets and regulatory network constructions. In contrast to benchmarking efforts that combine heterogeneous software environments or rely on manual parameter adjustment, our experimental design prioritizes uniform execution and standardized data handling across models.

The selected baseline methods reflect a broad range of modeling assumptions. GENIE3 [2] serves as a canonical expression-based method built on random forest regression and remains a strong classical reference point. GENELink [21] and GNE [56] leverage heterogeneous biological graphs and embedding strategies to encode regulatory structure beyond pairwise expression dependencies. STGRN [23] incorporates attention-based temporal modeling to capture dynamic regulatory patterns, while GNNLink [17] frames GRN inference as a deep graph link-prediction task. Together, these approaches represent the prevailing methodological directions in modern GRN inference research.

For all datasets and gene regulatory network variants, the baseline models were trained and evaluated using identical train, validation, and test edge splits. Hyperparameter selection was confined strictly to the validation split in each case, and test edges were fully withheld until final evaluation. GRNPred followed the same partitioning and tuning protocol, ensuring that observed differences in performance arise from representational and modeling differences rather than experimental artifacts.

To address the pronounced label imbalance intrinsic to GRN inference, we adopted a uniform bootstrap-based evaluation strategy across all methods. Each model was assessed over 100 bootstrap iterations, with all positive test edges retained and an equal number of negative edges sampled per iteration to form balanced evaluation subsets. Performance metrics based on the sampling strategy, including sampled AUROC and AUPRC, were computed for each replicate and reported as averages across all bootstrap rounds. This approach reduces variance due to different negative-edge sampling and yields more reliable and stable estimates of model performance.

## Data Availability

The datasets used in this study are publicly available. Original single-cell gene expression matrices and ground-truth regulatory networks were obtained from the BEELINE benchmark suite and can be accessed at https://zenodo.org/records/3701939. Gene Ontology annotations were retrieved from the Gene Ontology Consortium at http://purl.obolibrary.org/obo/go-basic.obo. Promoter sequences were extracted from the GRCh38.p14 human reference genome hosted by NCBI, available at https://www.ncbi.nlm.nih.gov/assembly/GCF_000001405.40/. Transcription start sites were defined using the GENCODE v48 gene annotation, obtained from https://ftp.ebi.ac.uk/pub/databases/gencode/Gencode_human/release_48/gencode.v48.annotation.gtf.gz. Position weight matrices were obtained from the JASPAR 2024 database, and gene text descriptions were retrieved from UniProtKB.

## Code Availability

The source code for GRNPred, including scripts for model training, evaluation, is publicly available at the GitHub repository: https://github.com/jianlin-cheng/GRNPred/tree/main.

## Author contributions

Conception: JC; Design of the method: JC, TN, and AH; Implementation and experimentation: TN and AH; Data collection and analysis: TN and AH; Manuscript writing: TN, JC, and AH.

## Acknowledgements

This work is supported in part by funds from the National Science Foundation (NSF grants: # 2343612, # 2308699, and #2525780) and from the Department of Energy (grant #: DE-SC0026121).

